# Evaluation of a Recessive Male Floret Color in Cultivated Northern Wild Rice (*Zizania palustris* L.) for Pollen-Mediated Gene Flow Studies

**DOI:** 10.1101/2021.01.26.428335

**Authors:** Clare Gietzel, Jacques Duquette, Lillian McGilp, Jennifer Kimball

## Abstract

Northern wild rice (NWR; *Zizania palustris* L.) is a wind-pollinated, annual, aquatic grass that grows naturally in the Great Lakes Region (GLR) of the United States and Canada, and is also cultivated in flooded paddies, predominantly in California and Minnesota. A better understanding of pollen-mediated gene flow is needed within the species for both conservation and breeding efforts as cultivation occurs within the species natural range and spatially-isolated, paddy structures are limited within breeding programs. Widely cited pollen travel research in NWR demonstrated that pollen could travel at least 3200m. However, a population segregating for male sterility was used as the pollen recipient in the study and was determined to not be adequate for NWR pollen travel studies. Here, we present the characterization of a recessive white male floret (WMF) population in contrast to the dominant, purple male floret (PMF) color of cultivated NWR along with estimates of pollen-mediated gene flow in a cultivated paddy setting. Studies conducted in 2018 and 2019 revealed that the primary amount of pollen-mediated gene flow occurred within the first 7m from the PMF donor source with no gene flow detected past 63m. These results suggest that the likelihood of pollen-mediated gene flow between cultivated NWR and natural stands remains low. We also identified a strong linkage between male floret, auricle, and culm color. This study demonstrates that the WMF trait is an excellent candidate for use in pollen-mediated gene flow studies in NWR.

## Introduction

Pollen-mediated gene flow in outcrossing species often represents the movement of unwanted alleles from one population to another. This movement can affect varietal purity and cause crop-to-wild or weedy relative gene flow (Ellstrand et al., 2003). The dispersion of pollen can depend on numerous factors including pollen viability, flowering synchrony, weather conditions, and surrounding vegetation (Shivanna et al., 1991; Gliddon et al., 1999; Thompson et al., 1999). In particular, weather conditions, such as wind speed, wind direction, and relative humidity, significantly affect pollen travel trends in wind-pollinated species (Latta and Mitton, 1999; Steinitz et al., 2011; Born et al., 2012). Despite this heavy environmental influence, most models predict the rapid dissipation of pollen densities following release from the source (Messeguer et al., 2006; Sarangi et al., 2017). For breeders and conservationists alike, an understanding of pollen travel trends for species of interest can aid in eliminating or reducing unwanted pollen-mediated gene flow.

The movement of pollen can be monitored using a wide range of techniques. Traps to track the physical movement and deposition of pollen, mostly commonly sedimentation and filtration traps, have been used across a range of species (Kearns and Inouye, 1993; Mullins and Emberlin, 1997). Other trapping methods involve the use of marked or tagged pollen (Moon et al., 2006). Specifically tracking pollen-mediated gene flow requires the use of morphological traits, such as kernel color (Hanson et al., 2005; Bannert and Stamp, 2007), cone size (Zobel, 1951), and leaf hair density (Hardig et al., 2000), or molecular markers, which can track specific genes (Ouborg et al., 1999). Molecular markers, in particular, have played an important role in tracking gene movement from genetically modified (GM) crops to non-GM varieties, landraces, and weedy relatives (Ma et al., 2004; Watrud et al., 2004; Pla et al., 2006).

Northern Wild Rice (NWR), *Zizania palustris* L., is an annual, wind-pollinated aquatic grass native to the Great Lakes Region of North America. It is a protogynous, monoecious species and its female florets are located above male florets on the panicle. Cultivation of the species began in Minnesota in the 1950s (Oelke et al., 1982) and has since become a high-valued specialty crop in major markets (Tuck, 2019). Breeding strategies and varietal development are largely based on phenotypic recurrent selection within open-pollinated populations (Grombacher et al., 1997). Spatially-isolated, permanent paddy structures are limited within breeding programs, necessitating a better understanding of pollen travel trends, especially within short distances, to maintain the genetic purity of breeding lines (Ireland et al., 2006; Baltazar et al., 2015). Additionally, cultivation occurs within the species’ center of origin, where the conservation of natural stands in lakes and streams is imperative to the maintenance of genetic diversity and delicate ecosystems. NWR is considered an indicator of overall ecosystem health, providing food and habitat for a variety of wildlife (Rogosin, 1954; Fannucchi, 1983). Therefore, an understanding of potential gene flow from natural stands to cultivated paddies and vice versa is an integral aspect of pollen travel research in NWR.

Initial pollen travel studies in NWR found diurnal pollen release patterns, similar to those in corn and other grass species, with the highest pollen concentrations occurring between 1200-1700 hours, 20-23°C, 50-60% relative humidity, and wind speeds of 7.5-9 kph (Cregan, 2004). Previous NWR pollen travel research also included the utilization of a bottlebrush (BB) population (Cregan, 2004), with a compact male flower morphology, linked to an uncharacterized male sterility gene (Stucker et al., 1984; Grombacher et al., 1997). Cregan (2004) reported that NWR pollen could travel at least 3200m, a number that has been widely cited as validation of concerns regarding pollen-mediated gene flow in NWR. However, the BB trait is considered inadequate for NWR pollen travel studies due to the somewhat weak linkage between the BB trait and male sterility, leading to pollen-producing BB plants and resulting seed set (contamination). In this study, we compare two populations of cultivated NWR, with either a recessive white male floret (WMF) or a dominant, purple male floret (PMF), and use them to estimate pollen-mediated gene flow in a cultivated paddy setting. This study demonstrates that the WMF trait is an excellent candidate for use in pollen-mediated gene flow studies in NWR.

## Materials and Methods

### Plant Material

In this study, a homozygous recessive WMF population (Figure 1a) was utilized to estimate pollen-mediated gene flow in NWR. Multiple PMF populations (Figure 1b) were utilized as pollen donors to maximize the flowering synchronicity between WMF and PMF populations. The 2018 flowering period for the WMF population was from July 15^th^ to August 15^th^ and from July 25^th^ to August 25^th^ in 2019.

**Figure 1.**
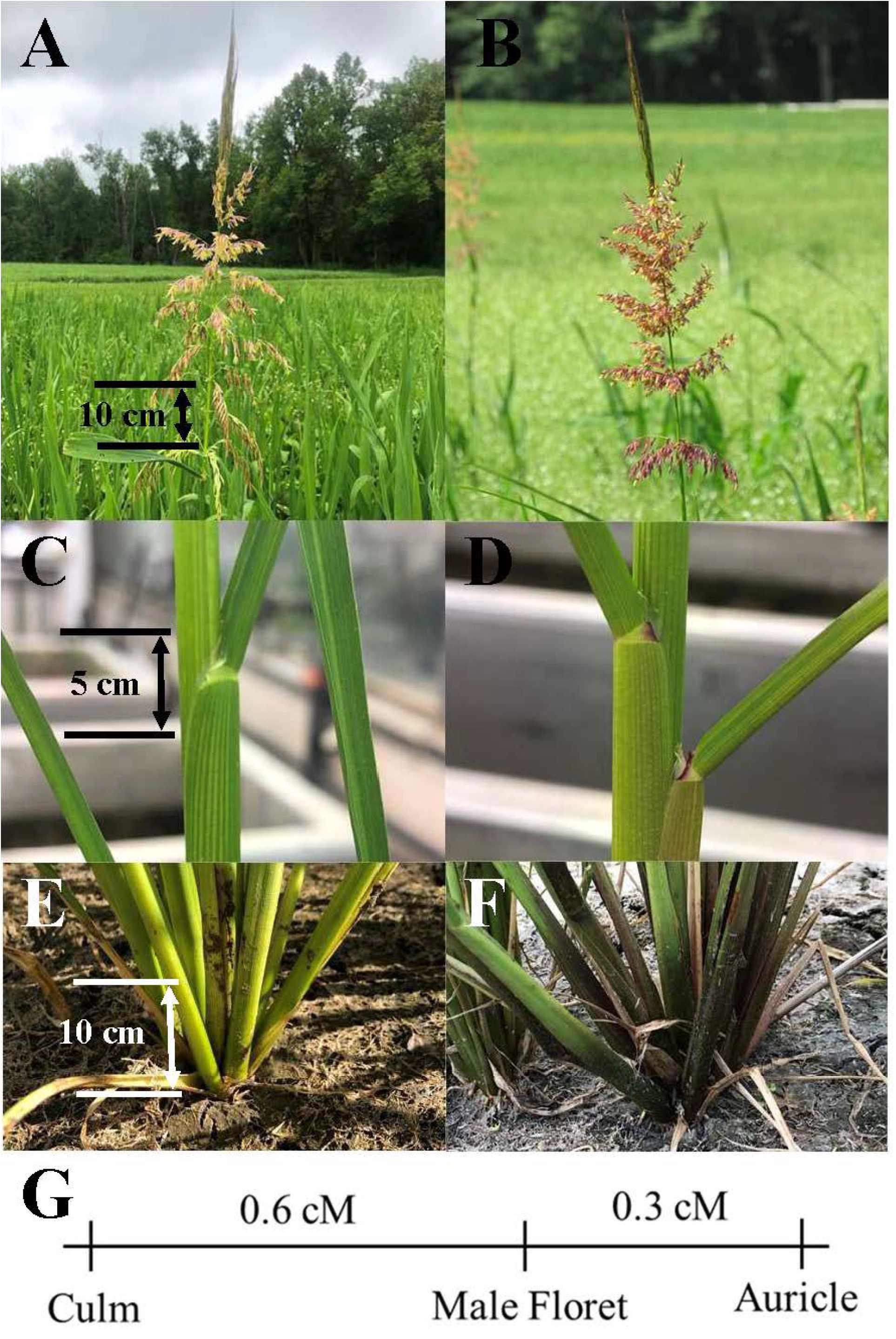
Phenotypes of the purple male floret (PMF) and white male floret (WMF) colors in northern wild rice (NWR). The left side of the figure represents white phenotypes while the right side represents purple phenotypes. A/B, C/D, and E/F represent the male floret, auricle, and culm, respectively. Finally, G represents the putative recombination map for the genes associated with these traits.

### Weather Data

Wind speed (km/h) and direction data were collected from the National Oceanic and Atmospheric Administration (NOAA) weather station located at the Itasca County Airport in Grand Rapids, MN, approximately 3.3 miles from the trial. Air temperature (°C) and accumulated precipitation (mm) were collected from an automated weather station at the Grand Rapids U.S. Forest Research Service Lab in Grand Rapids, MN, approximately 0.5 miles from the trial.

### Pollen Travel Experiments

Experiments were conducted at the University of Minnesota North Central Research and Outreach Center (NCROC) in Grand Rapids, Minnesota (47.2372° N, 93.5302° W, and 392 m elevation) at the cultivated NWR paddy complex, where individual paddies have 1-1.5m raised dikes for irrigation purposes. In 2018, twenty 3m x 6m WMF plots with 3m single rows spaced 0.38m apart and 1m alleys were planted at 15.7 kg ha^−1^. These plots were planted 3m south/southeast of a 24-plot (12m x 41m) PMF trial, which served as the purple pollen donor (Figure 2a). At its farthest distance, a WMF plot was a maximum of 35m from a PMF plot (north to south). In 2019, the experiment was expanded to include larger pollen travel distances and forty-four 3m x 6m WMF plots were planted with the northwestern-most plot consisting of multiple PMF genotypes (Figure 2b). The maximum distance between a WMF plot and the PMF plot was 9m, west-east, and 70m, north-south. The experimental design for these experiments was largely dictated by the size and shape of paddies (~15-25m wide and ~90m long), which eliminated the possibility of planting a PMF plot in the middle of a large WMF plot to evaluate 360 degrees of pollen travel from the source. Instead, common summer wind patterns were utilized to establish dominant wind directions at the paddy complex, resulting in the PMF donor source being planted directly north of the WMF plots.

**Figure 2.**
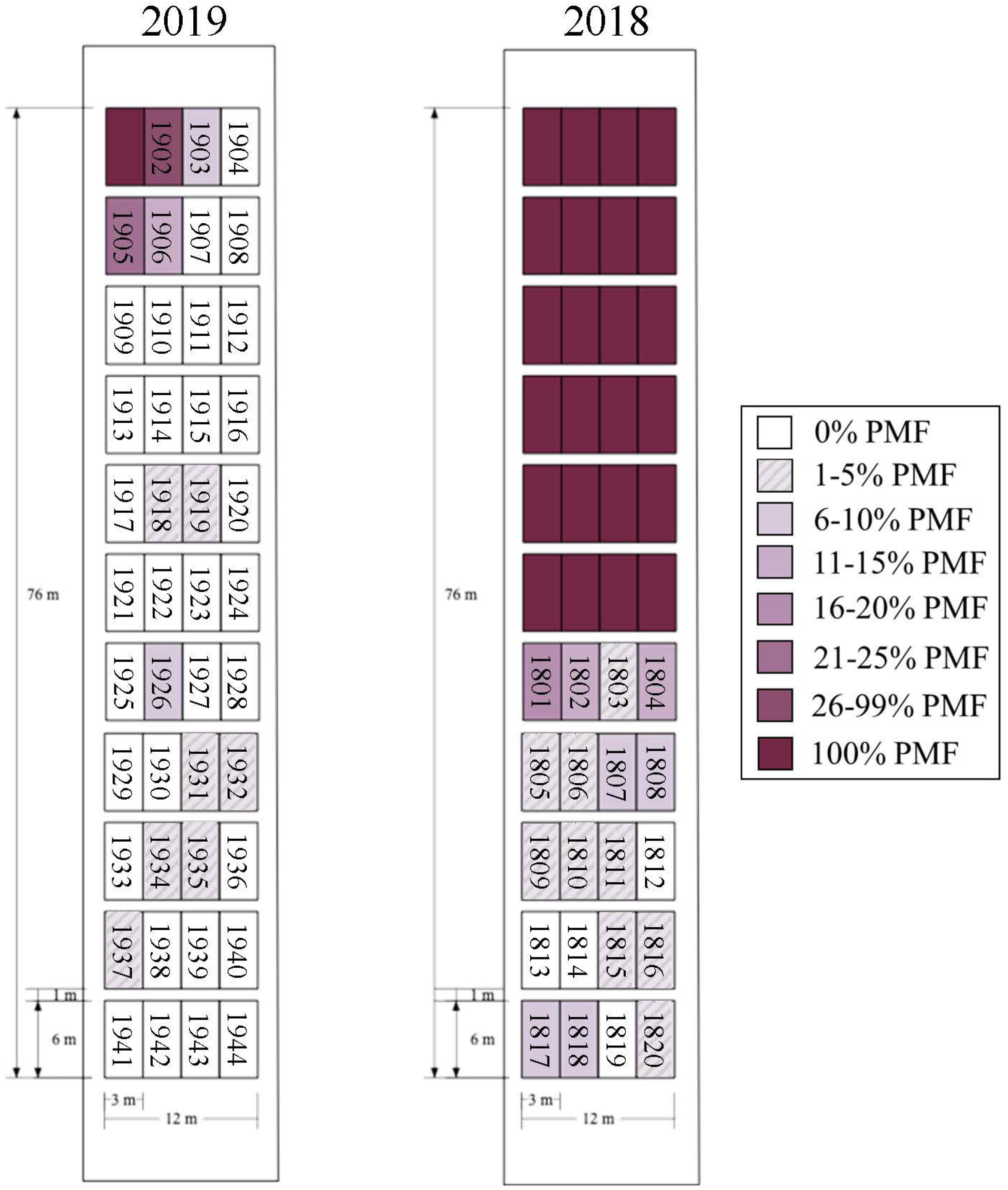
Plot maps of 2018-2019 pollen travel experiments in northern wild rice (NWR) along with progeny testing results, which indicate purple male floret (PMF) hybridization rates in the white male floret (WMF) progeny. White boxes represent 100% WMF plots and purple boxes represent varying degrees of PMF densities.

Paddies were amended with 16.7 kg ha^−1^ Environmentally Safe Nitrogen (ESN), 9.3 kg ha^−1^ urea, and 7.4 kg ha^−1^ potassium. Immediately following planting, paddies were flooded to 20cm average water depth. Copper sulfate (1.13 kg ha^−1^ rate) was used to control algal growth and Aquabac (200G) (*Bacillus thuringiensis* subsp. *israelensis*-Becker Microbial Products, Coral Springs FL; 1.86 kg ha^−1^ rate) to control aquatic pests, specifically midge larvae. Plots were monitored daily for PMF plants in the WMF plots, which were rogued prior to pollen set. Individual WMF plots were harvested each year, processed, and stored in the dark, at 3°C, on water, for the duration of the seed dormancy period.

### Progeny Testing

For both years of the study, ~50-100 stratified seeds per WMF plot were germinated on petri dishes lined with filter paper and hydrated with 10 ml of H_2_O. The petri plates were placed at an ambient temperature of ~23°C, under LabLink LED LabLights (PG LifeLink, Erlanger, KY), set to a 16-hr day length. After 10 days, 5cm long seedlings were transplanted into individual 25cm deep plant cones, submerged in 680L aquaponics tanks. The soil in each cone was amended with 22.4 kg ha^−1^ N, in the form of urea, and 0.125 g iron chelate. Throughout growth and flowering, plants were monitored daily for culm, auricle, and floret color. Anecdotal evidence within the program suggests a potential link between the genes controlling these traits. In the spring of 2019, ~38 progeny plants per WMF plot, from the 2018 study, were evaluated. For the 2019 trial, only ~22 progeny plants per WMF plot were evaluated in the spring of 2020 due to the COVID-19 global pandemic.

### Data Analysis

Monthly mean air temperature was calculated by averaging the daily maximum and minimum air temperature values. Cumulative precipitation was calculated by adding together the daily precipitation, in millimeters, for all days in a given time period. Analysis of wind speed and direction data was conducted using R 4.0.1 (R Core Team, 2020), and the OpenAir package (Carslaw & Ropkins, 2012) was used to generate wind roses with the use of ggplot2 and ggplotify (Wickham, 2016; Yu, 2020). Wind run, or the total distance that the wind traveled over a given timeframe, was calculated on a 24 hr basis by multiplying the average wind speed by the amount of time elapsed per data point. Data was collected hourly unless a shift in wind direction occurred, in which case, another data point was generated. In order to analyze differences in wind run per plot, the degree range for each of the 16 cardinal directions was superimposed onto each plot map and wind directions that conferred a high probability of wind traveling through pollen donor PMF plots to each WMF plot was determined. An empirical model of pollen-mediated gene flow was obtained by regressing the hybridization rate on the maximum distance from the PMF pollen source using the NLIN procedure in SAS statistical software package v9.4 (SAS Institute, 2013) similar to Schmidt *et al*. (2013). The negative exponential model: hybridization rate = α^−β(distance)^ was used. The suitability of the model fit was verified using regression coefficients.

Percent hybridization per plot was calculated as the proportion of WMF plot progeny with purple male florets over the total progeny tested. To evaluate the effect of distance from the PMF source, the distance in meters from the closest edge of the PMF plot(s) to the furthest edges of each WMF plot was measured and then pooled for each range of the experiment. Recombination frequencies between culm, auricle, and male floret color were calculated using parental (all white or all purple phenotypes) and recombinant class data from 2018. The number of plants in each recombinant class divided by the total number of observed parental phenotypes provided estimates of genetic linkage in centiMorgans (cM) between the three traits.

## Results and Discussion

### Weather Data

Throughout the growing period in 2018, the mean air temperature (°C) was generally warmer than the 20-year average for Grand Rapids, Minnesota but in 2019, it was cooler than the 20-year average (Table 1). Temperature differences between 2018 and 2019 were minimal, with the exception of May and August, when it was more than 2°C cooler in 2019. During the 2018 and 2019 flowering periods (July 15^th^-August 15^th^ and July 25^th^-August25^th^, respectively), weekly mean air temperatures varied widely. The last week of July and first week of August were peak flowering weeks during both years and had a mean air temperature of 20.00°C and 17.98°C in 2018, and 13.89°C and 19.1°C in 2019, respectively. In other Poaceae species, temperature is known to affect anther ripening, where warmer weather can confer higher concentrations of shed pollen (Hart et al., 1994), while low temperatures can reduce meiotic efficiency during anthesis (Zeng et al., 2017). Cregan (2004) found NWR pollen shed to be highest between 20-23°C, which suggests warm, but not hot, temperatures improve pollen shed. It is possible that the low temperatures during the first week of peak flowering in 2019 decreased pollen shed. However, more research regarding the relationship between temperature and pollen shed in NWR is needed to validate this conclusion.

**Table 1.** Weather data including mean air temperature (°C) and precipitation (mm) during the growing season of 2018 and 2019 in Grand Rapids, Minnesota. Yearly precipitation values are totaled per month and cumulative precipitation values (in parentheses) give the running total for the entire growing season.

| Month | PGS<sup>†</sup> | Mean Air Temperature (°C) |  |  | Monthly Precipitation (mm) |  |  |
| --- | --- | --- | --- | --- | --- | --- | --- |
|  |  | 2018 | 2019 | 20 Year Average | 2018 | 2019 | 20 Year Average |
| May | 0 | 15.16 | 9.44 | 11.92 | 95.50 (95.50) | 45.72 (45.72) | 87.00 |
| June | 1-2 | 17.83 | 16.81 | 17.35 | 119.38 (214.88) | 151.13 (196.85) | 115.06 |
| July | 2-5 | 20.39 | 20.38 | 20.50 | 82.55 (297.43) | 61.21 (258.06) | 84.84 |
| August | 5-8 | 19.07 | 16.30 | 18.88 | 35.05 (332.49) | 61.47 (319.53) | 79.86 |
| September | 7-9 | 13.72 | 14.52 | 14.36 | 80.26 (412.75) | 117.35 (436.88) | 81.56 |

<sup>†</sup>Duquette & Kimball, 2020

Monthly precipitation (mm) in 2018 was similar to the 20 year average with the exception of the month of August, which was ~43% drier than the 20 year average (Table 1). In 2019, monthly precipitation levels were highly variable from month to month during the growing season and their 20 year averages. May, July, and August were drier in comparison, while June and September were wetter. When comparing 2018 and 2019, accumulation varied widely between the months. During the flowering periods, plots received a total of 12.96 mm of precipitation in 2018 and 78.22 mm in 2019. While the effect of precipitation on pollen travel in NWR has not been previously explored, studies have shown that pollen concentration in the air decreases significantly on rainy days, as rain washes pollen grains from the air (Scott, 1970; Hart et al., 1994). In 2018, there were 11 days during the 30-day flowering period with rainfall, however only 3 of those days accumulated to more than 1 mm. In 2019, there were 17 days of recorded rainfall during the 30-day flowering period but only 11 with more than 1 mm.

Wind direction and speed are primary drivers of pollen travel in wind-pollinated species. In 2018, winds originated predominantly from the northwest and southeast, while winds in 2019 primarily arose from the south (Figure 3). In 2018, ~33% of the wind came from the northwest (NW, NNW, WNW), and ~31% came from the south (S, SSE, SSW). In 2019, ~23% of the wind came from the northwest, while ~46% came from the south (Figure 3).). Mean wind speeds were similar between years with speeds reaching 7.9 km/h in 2018, and 8.2 km/h in 2019. Both years had similar amounts of calm air during the flowering periods (38.4% and 36.4%, respectively). Both years had similar levels of calm air during the flowering periods (38.4% and 36.4%, respectively). Both years also had similar frequencies of high speed wind. During the flowering period in both years, approximately 1% of the wind that blew was characterized as a “fresh breeze,” defined as 29-38 km/h in the Beaufort scale. In 2018, 7% of the wind that blew was considered a moderate breeze (20-28 km/h), compared with 8% in 2019. There were more days during the 2019 flowering period with recorded wind gusts (20) than in 2018 (16). There is scientific evidence to suggest that a slight variation in wind speed, irrespective of an increase or decrease, has a greater effect on pollen travel than higher wind speed alone (Sauliene and Veriankaite, 2012) and therefore, these wind gusts may play an important role in moving NWR pollen from the pollen donor source.

**Figure 3.**
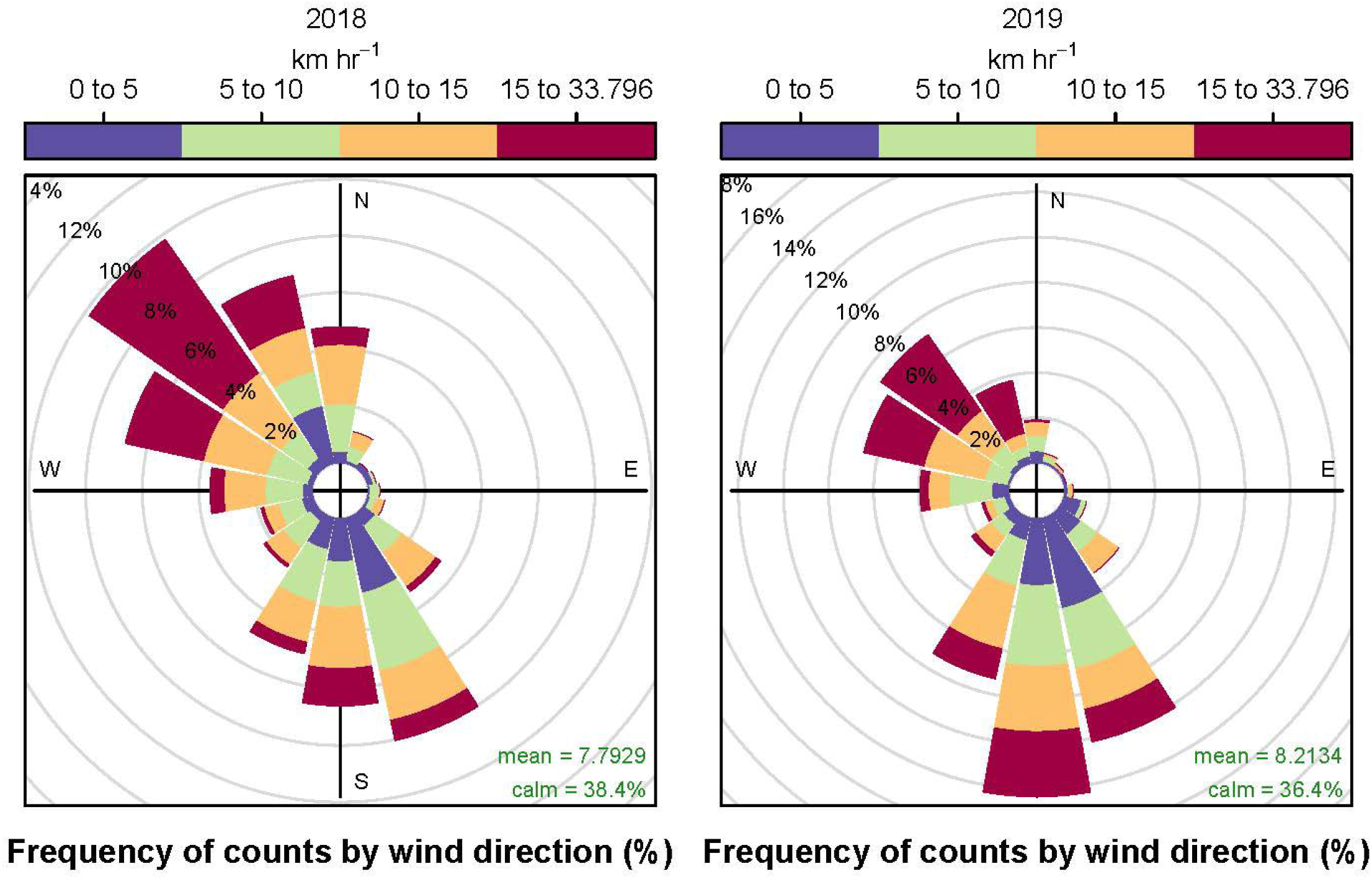
Wind trends during the 2018 and 2019 flowering periods at the Itasca County Airport in Grand Rapids, MN. Wind trends show wind blowing toward the direction specified.

### Distance and Direction of Pollen Travel

Detection of PMF pollen-mediated gene flow in WMF plots ranged from 0-19% and 0-48% in 2018 and 2019, respectively (Figure 2). In both years, the highest rate of PMF penetrance occurred within the WMF plots bordering or closest to the PMF donor plot(s). This suggests that the highest percentage of pollen-mediated gene flow within NWR occurs within the first 7m of the pollen donor source. Beyond a 7m N-S distance, pollen-mediated gene flow from PMF plots to WMF plots ranged from 1.24%-5.32% in 2018 (up to 35m tested) and 0.00-2.44% in 2019 (up to 70m tested) (Table 2). No gene flow was detected past 63m. However, it is important to note that pollen density would be higher in a larger production field, which could result in higher rates of hybridization. The higher rates of pollen penetrance into the 2018 plots may have been influenced by the amount of donor pollen available per the experimental design, the higher mean air temperatures, or the lower amounts of precipitation in 2018. While we are unable to confirm using this dataset, we hypothesize that PMF pollen penetrance in 2019 beyond 49m was caused by contaminant PMF plants that were not identified and rogued in WMF plots prior to pollen shed. These contaminant plants were identified in plots 1934 and 1935. However, further studies need to be conducted to confirm this. Despite this possibility, the estimates of pollen-mediated gene flow in this study were comparable to those identified in other grass species. In both maize (*Zea mays*) (Messeguer et al., 2006) and common waterhemp *(Amaranthus rudis* Sauer) (Sarangi et al., 2017), evidence of pollen-mediated gene flow via wind was found within the first 50m surrounding pollen donor plants but then dropped off considerably.

**Table 2.** Effect of north-south (N-S) distance from the pollen donor source on pollen-mediated gene flow from purple male floret (PMF) northern wild rice plots to white male floret (WMF) plots in Grand Rapids, Minnesota during the 2018 and 2019 seasons.

|  | **<u>Hybridization rate (0.0093SE)</u>** |  |
| --- | --- | --- |
| **N-S Distance (m)** | **2018** | **2019** |
| 1-7 | 13.07% | 9.20% |
| 8-14 | 5.32% | 0.00% |
| 15-21 | 1.74% | 0.00% |
| 22-28 | 1.24% | 2.38% |
| 29-35 | 3.92% | 0.00% |
| 36-42 | - | 2.44% |
| 43-49 | - | 2.41% |
| 50-56 | - | 2.35% |
| 57-63 | - | 1.22% |
| 64-70 | - | 0.00% |

To investigate the relationship between the rate of hybridization in WMF plots and distance from the purple pollen donor source among years, we fit a curve to the data from each year and compared the confidence intervals for α and β (Table 3). The model was deemed appropriate for the description of pollen-mediated gene flow in both years by the highly significant parameter estimates and the non-zero confidence intervals. However, this empirical model only includes distance as a predictive factor of pollen-mediated gene flow and does not account for other environmental factors, such as wind. Therefore, we generated an ANOVA using a multiple regression model including distance and wind run. Both variables had a significant effect in both years during the flowering periods. In this regression model, distance had the biggest impact on the rate of hybridization in 2018, while wind run had a bigger impact on hybridization in 2019 (Table 4). The significant interaction between distance and wind run indicates that distance from the pollen source as well as wind speed and its proportion have an impact on hybridization. These results suggest that as distance from the pollen source increases, wind becomes a more important factor in the rate of hybridization, a phenomenon that has been identified in other species as well (Hanson et al., 2005; Schmidt et al., 2013). While we suggest the possibility that contamination could have led to hybridization at greater distances, it is also possible that wind speed and direction could have led to hybridization from the purple pollen source.

**Table 3.** Analysis of variance and parameter estimates obtained by fitting hybridization frequency to distance from the source population using the model: frequency of hybridization = α exp(–β × distance).

| Year | Source | df | MSE<sup>†</sup> | P>F | Coefficient | Estimate | SE | Confidence Interval |  |
| --- | --- | --- | --- | --- | --- | --- | --- | --- | --- |
| **2018** | Model | 2 | 0.0427 | <0.0001 | $\alpha$ | 0.2524 | 0.0272 | 0.1952 | 0.3096 |
| | Error | 18 | 0.0009 | | $\beta$ | 0.0199 | 0.0016 | 0.0166 | 0.0232 |
| **2019** | Model | 2 | 0.0540 | 0.0003 | $\alpha$ | 0.1256 | 0.0282 | 0.0686 | 0.1826 |
| | Error | 41 | 0.0055 | | $\beta$ | 0.0076 | 0.0014 | 0.0048 | 0.0104 |

<sup>†</sup>Mean square error

**Table 4.** Analysis of variance tests for the effects of distance and wind run on the frequency of hybridization in 2018 and 2019.

| Year |  | df | F value | P value |
| --- | --- | --- | --- | --- |
| 2018 | Distance | 1 | 18.916 | 0.0005 |
|  | Wind Run | 1 | 5.690 | 0.0298 |
|  | Distance * Wind Run | 1 | 9.795 | 0.0065 |
| 2019 | Distance | 1 | 10.811 | 0.0021 |
|  | Wind Run | 1 | 31.505 | 0.0002 |
|  | Distance * Wind Run | 1 | 7.997 | 0.0074 |

### Recombination between Culm, Auricle, and Male Floret Color

In NWR, purple and white color variation also presents in the culm (Figure 1c-d) and auricle (Figure 1e-f), which are hypothesized to be linked to male floret color. To assess the degree of linkage, observations of floret, culm and auricle color were collected in 2018. Of the 335 analyzed plants, only three plants displayed recombinant genotypes between male floret color, culm color, and auricle color with an estimated 0.9 cM separating culm and auricle color genes (Figure 1g). The limited recombination between these three traits indicates they are tightly linked, which will be helpful for progeny testing in future WMF travel studies, as culm and auricle color can be evaluated at earlier stages of development. Genes for purple coloration (anthocyanin accumulation) in various plant parts have been identified in other grain crops including wheat, triticale, and white rice (Dhulappanavar, 1973; Dhulappanavar, 1979; Zeven, 1985).

## Conclusions

This study confirmed that the utilization of the male floret color as a means to track pollen-mediated gene flow in NWR is an effective strategy, as long as the WMF seed source is not contaminated with seed harboring the purple male floret color. The contamination in the WMF seed sources highlights the difficulties of controlling pollen travel in cultivated NWR breeding programs. However, pollen-mediated gene flow in this study was limited beyond 7m from the pollen donor source, providing estimates of pollen-mediated gene flow for establishing new pollen travel mitigation strategies in irrigated paddy settings. The limited range of pollen-mediated gene flow in NWR is further supported by genetic diversity studies that have estimated high genetic differentiation between different lake and river populations of NWR, suggesting limited gene flow among populations (Lu et al., 2005; Xu et al., 2015). These results suggest that the possibility of pollen-mediated gene flow between cultivated NWR and natural stands remains low. However, Halsey et al. (2005) indicated that the relative amount of pollen produced from the pollen source is an important factor when evaluating pollen-mediated gene flow in species. Our study was small in scale in comparison to cultivated paddies as well as natural stands in lakes and river systems. In the future, we suggest a large-scale on-farm study, similar to Cregan (2004), is necessary to effectively determine maximum pollen travel distances in NWR. Additionally, while we identified that air temperature, rainfall, and wind patterns play a role in the distance of pollen travel in NWR, more research is needed to fully understand the effect of weather conditions on pollen travel.

